# Clock-dependent chromatin accessibility rhythms regulate circadian transcription

**DOI:** 10.1101/2023.08.15.553315

**Authors:** Ye Yuan, Qianqian Chen, Margarita Brovkina, E. Josephine Clowney, Swathi Yadlapalli

## Abstract

Chromatin organization plays a crucial role in gene regulation by controlling the accessibility of DNA to transcription machinery. While significant progress has been made in understanding the regulatory role of clock proteins in circadian rhythms, how chromatin organization affects circadian rhythms remains poorly understood. Here, we employed ATAC-seq (Assay for Transposase-Accessible Chromatin with Sequencing) on FAC-sorted Drosophila clock neurons to assess genome-wide chromatin accessibility over the circadian cycle. We observed significant circadian oscillations in chromatin accessibility at promoter and enhancer regions of hundreds of genes, with enhanced accessibility either at dusk or dawn, which correlated with their peak transcriptional activity. Notably, genes with enhanced accessibility at dusk were enriched with E-box motifs, while those more accessible at dawn were enriched with VRI/PDP1-box motifs, indicating that they are regulated by the core circadian feedback loops, PER/CLK and VRI/PDP1, respectively. Further, we observed a complete loss of chromatin accessibility rhythms in *per^01^* null mutants, with chromatin consistently accessible throughout the circadian cycle, underscoring the critical role of Period protein in driving chromatin compaction during the repression phase. Together, this study demonstrates the significant role of chromatin organization in circadian regulation, revealing how the interplay between clock proteins and chromatin structure orchestrates the precise timing of biological processes throughout the day. This work further implies that variations in chromatin accessibility might play a central role in the generation of diverse circadian gene expression patterns in clock neurons.

**Significance Statement:** Chromatin organization plays a critical role in gene regulation in development and in disease. In this study, we discovered robust circadian oscillations in the chromatin accessibility of regulatory elements of clock-regulated genes in *Drosophila* clock neurons, with enhanced accessibility either at dusk or dawn, which correlated with their peak transcriptional activity. We found enrichment of E-box motifs in genes that exhibited enhanced accessibility at dusk, and enrichment of VRI/PDP1-box motifs in genes that exhibited higher accessibility at dawn. Moreover, the complete loss of chromatin accessibility rhythms in *per^01^* mutants highlights the essential role of the Period protein in driving chromatin compaction during the repression phase. This study highlights the significance of chromatin organization in the generation of ∼24-hour circadian rhythms.

## Introduction

Chromatin organization plays a pivotal role in ensuring proper gene expression patterns during development, differentiation, and in response to environmental stimuli (1). Chromatin can adopt either an “open” or “closed” configuration. Euchromatin is less condensed (open) and is associated with active gene transcription, while heterochromatin is more compact (closed), which makes the DNA less accessible, and is generally associated with gene repression (2). For example, during the developmental transition from a stem cell to a more specialized cell type, the promoters, enhancers, and other regulatory elements of specific genes necessary for the function of the differentiated cell become more accessible to transcriptional machinery, which enables their expression (3, 4). Conversely, other regions become less accessible, leading to gene silencing. For instance, during mammalian development, key developmental genes like the Hox loci undergo a significant change in chromatin structure, becoming highly accessible during the differentiation process (5).

While the essential role of chromatin accessibility in controlling gene expression during development is widely recognized, much less is known about chromatin accessibility dynamics over the 24-hour circadian cycle and its role in driving circadian rhythms. Circadian clocks are cell-autonomous timekeepers that orchestrate ∼24-hour rhythms in the expression of a large number of genes, thereby controlling much of our physiology and behavior, including sleep-wake cycles, and metabolism (6–8). Prior studies have demonstrated that the core clock transcription machinery recruits a variety of epigenetic remodelers—including histone-modifying enzymes, Polycomb complexes, and ATP-dependent chromatin remodeling enzymes—to orchestrate rhythmic gene expression over the circadian cycle (8). Further, previous work has identified rhythmic patterns in promoter-enhancer contacts of clock genes over the circadian cycle in mouse liver and kidney cells (9, 10). Additionally, both our past work (11) and that of others (12) have shown that subnuclear location of clock-regulated genes at the nuclear periphery is critical for their rhythmic gene expression. However, what remains less well understood is the broader dynamics of chromatin accessibility within clock neurons and its impact on circadian rhythms.

Circadian transcription could predominantly be governed by specific protein factors, while the underlying chromatin structure remains stable over the 24-hour circadian cycle. However, an alternative hypothesis is that the clock might actively modulate chromatin accessibility rhythms, thereby producing daily transcriptional oscillations. To systematically explore these possibilities, we utilized *Drosophila melanogaster* as it has a well conserved, yet relatively simple circadian clock system. The *Drosophila* clock network consists of ∼150 clock neurons that express clock proteins (13, 14) (**Figure S1A**). Circadian clocks in all eukaryotes are based on negative transcription-translation delayed feedback loops (6). In *Drosophila*, the circadian clock is primarily regulated by two key feedback loops involving the PERIOD/CLOCK and VRILLE/PDP1 proteins (6). CLOCK (CLK) and CYCLE (CYC) proteins serve as transcriptional activators, binding to the E-boxes (enhancer-box) of clock-regulated genes, including core clock genes such as *period* (*per*), *timeless* (tim), *vrille* (vri), and *pdp1-ε*, and activating their transcription around dusk (**Figure 1A**). After a time-delay, PER and TIM proteins enter the nucleus around dawn, where they function as transcriptional repressors. There, they counteract the activities of CLK/CYC, inhibiting the transcription of both their own genes and other clock-regulated genes. The sequential binding of CLK/CYC activators to chromatin around dusk followed by PER/TIM repressors around dawn orchestrates rhythmic gene expression patterns (**Figure 1A**).

**Figure 1.**
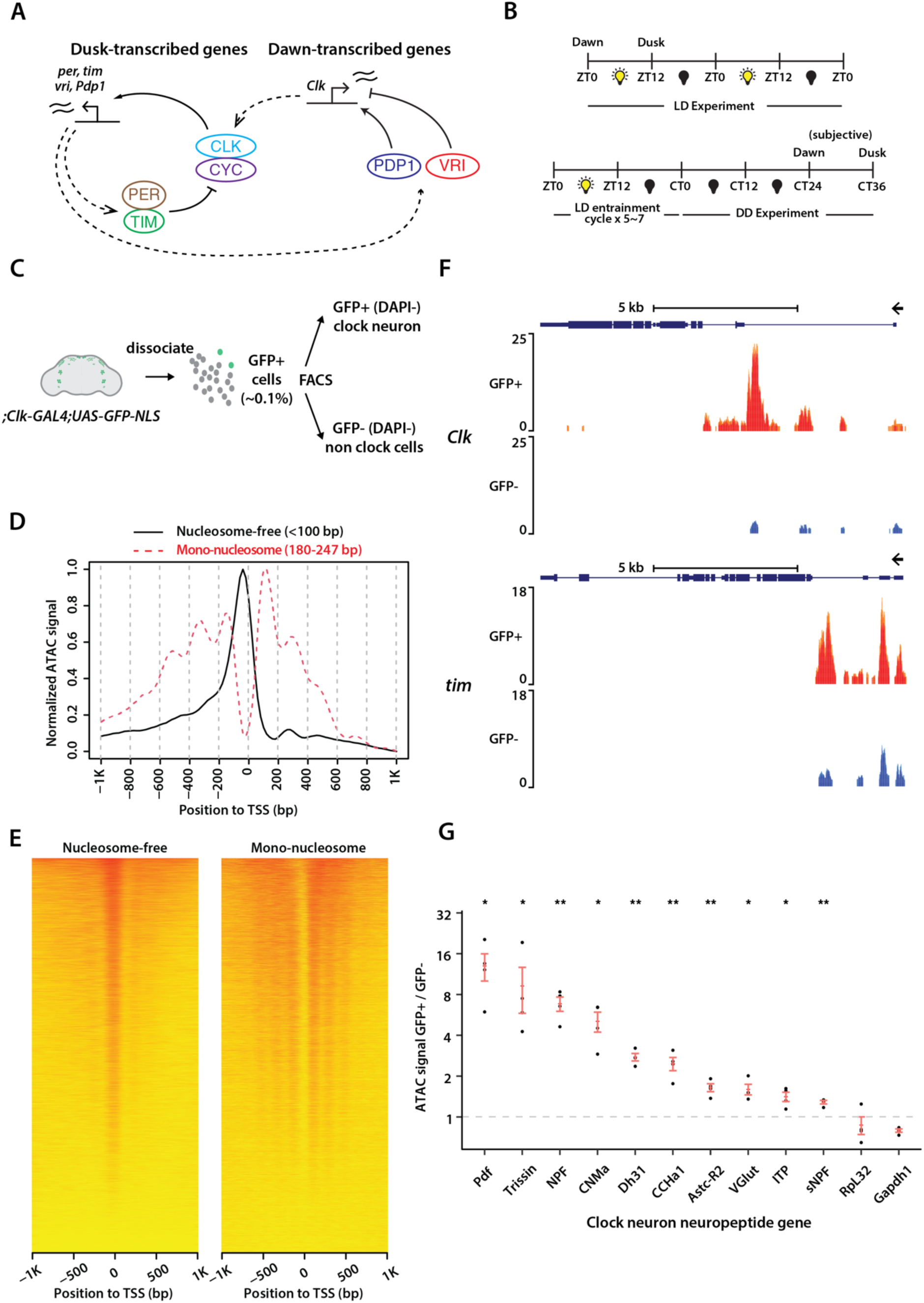
ATAC-sequencing of *Drosophila* clock neurons over the circadian cycle. (**A**) Schema of the *Drosophila* molecular clock feedback loops. (**B**) Experimental schema. Flies were entrained for 5 days in light-dark cycles (ZT0: lights-on/dawn, ZT12: lights-off/dusk) for LD experiments and then optionally released into complete darkness for DD experiments (CT0/CT12: subjective dawn/dusk). (**C**) Overview of clock neuron isolation procedure. Nuclear GFP signal is expressed with *Clk-GAL4* driver and cell suspension was prepared from ∼60 brains for each sample. Fluorescence activated cell sorting is used to sort live clock neurons with DAPI as viability marker. GFP-negative cells are sorted as non-clock-neuron control. (**D**) Representative ATAC-signal distribution aligned to transcription start sites (TSS) as reported by ATACseqQC. Fragments are classified into different groups according to alignment length, including the nucleosome-free (<100bp) and mono-nucleosome (180∼247bp) groups. Clear TSS enrichment of nucleosome-free signal and nucleosome ladder are observed. (**E**) Heatmap of normalized ATAC signal distribution aligned to TSS. (**F**) Representative ATAC signal pile-up tracks at *Clock* and *timeless* loci. Signal is normalized to total number of reads for each sample and therefore is comparable among samples. Four different replicates are overlaid. Clock neurons (GFP+, DAPI-) show strong ATAC signal in *Clk* and *tim* loci while non-clock cells (GFP-, DAPI-) show significantly lower readout. (**G**) Comparison of ATAC signal at various clock-neuron neuropeptide gene loci between clock-neurons and non-clock cells. While housekeeping genes (*RpL32* and *Gapdh1*) does not show clock-neuron enrichment, all clock-neuron neuropeptide genes are significantly more accessible in clock neurons. The statistical test used was a two-sided Student’s t-test. *P < 0.01, **P < 0.001.

The VRI/PDP1 feedback loop forms the second key regulatory circuit. Around dusk, VRI, acting as a repressor, binds to specific sites (VP-box) within the *Clk* enhancer to inhibit its transcription. Conversely, at dawn, PDP1-ε, acting as an activator, binds to these VP-boxes within the *Clk* enhancer to promote its transcription (**Figure 1A**). This alternating binding activity of PDP1-ε (activator) and VRI (repressor) results in the rhythmic expression of *Clk* mRNA (15). This VRI/PDP1 feedback loop also plays a crucial role in controlling the rhythmic expression of numerous clock output genes, thereby influencing rhythmic behavior (16, 17). These two interlinked feedback loops, PER/CLK and VRI/PDP1, generate gene expression rhythms with opposite phases, where genes regulated by the PER/CLK loop peak around dusk while those regulated by the VRI/PDP1 loop peak around dawn (**Figure 1A**), thereby helping establish a robust circadian rhythm in *Drosophila*.

To investigate the impact of chromatin dynamics on circadian rhythms, here, we adapted ATAC-seq (18) (Assay for Transposase-Accessible Chromatin) to assay chromatin accessibility genome-wide from FAC (Fluorescence-activated cell)-sorted *Drosophila* clock neurons at multiple timepoints over the circadian cycle. Our studies revealed that circadian transcriptional activation and repression are accompanied by dynamic chromatin accessibility states that poise the genome for rhythmic gene expression. Specifically, we found that regulatory elements of clock-regulated genes undergo diurnal oscillations in chromatin accessibility over the circadian cycle. Motif analysis identified that genes that were more accessible around dusk contained E-boxes, while those that were more accessible around dawn contained VP-boxes. We also found that chromatin accessibility rhythms in clock neurons are completely abolished in arrhythmic *per^01^*null mutants (19). In summary, these findings highlight the important role of chromatin organization in circadian regulation, demonstrating how the interplay between chromatin accessibility and clock proteins affects circadian rhythms.

## Results

### ATAC-sequencing of *Drosophila* clock neurons over the circadian cycle

To investigate genome-wide chromatin accessibility within *Drosophila’s* clock neurons, we performed transposase-accessible chromatin sequencing (ATAC-seq) (18), a strategy that enables identification of accessible regions within the chromatin. To isolate and target clock neurons specifically, we used the *Clk856-GAL4* driver (20)—which is expressed in almost all Drosophila clock neurons (21)—to express EGFP in their nuclei (Fig. 1B). The *Clk856-GAL4>EGFP-NLS* flies displayed normal locomotor-activity rhythms in both light-dark cycles (LD12:12) and constant darkness (DD), displaying ∼24-hour period rhythms and activity peaks around dawn and dusk (**Figure S1B**). To conduct our ATAC-seq experiments, we entrained these flies to a 12:12 LD cycle, and then released them into constant darkness (DD) conditions. Flies were collected either at dawn (ZT0) or dusk (ZT12) in LD, or at subjective dawn (CT24) or subjective dusk (CT36) on the second day of DD (**Figure 1B**). Subsequently, we dissected fly brains, isolated clock neurons using fluorescence-activated cell sorting (FACS), separated nuclei from cytoplasmic content, performed TN5 transposition reaction, and prepared ATAC-seq libraries for paired-end sequencing as described previously (22) (**Figure 1C**). We performed four biological replicates for each condition and timepoint. For each of our experiments, we dissected ∼60 brains from both male and female flies which yielded ∼2,000 GFP-positive clock neurons. Additionally, we collected ∼10,000 GFP-negative cells, representing non-clock neurons which comprise a diverse set of neurons and glia in the brain, to serve as our control group. GFP-positive clock neurons constituted 0.1% of cells extracted from fly brains, consistent with previous estimates (21) (**Figures S1C, S1D**).

After mapping the sequencing reads to the *D. mel* genome (*dm6*) and filtering out PCR duplicates that were a by-product of the library amplification process, we were able to achieve a median depth of ∼5-10 million unique, high-quality mapped reads per sample. The high Pearson correlation between the biological replicates for each condition (a minimum of 0.94 across all samples) attests to the high reproducibility of our sequencing results and the notable similarity in chromatin profiles across all conditions (**Figure 2B**). Analysis of ATAC-seq signal across gene units from both LD and DD conditions revealed enrichment near the transcriptional start sites (TSSs), denoting the accessible chromatin within promoter regions (**Figures 1D, 1E, S1E**). We deposited all the ATAC signal pile-up tracks on UCSC Genome Browser (https://tinyurl.com/2ftuhv5p).

**Figure 2.**
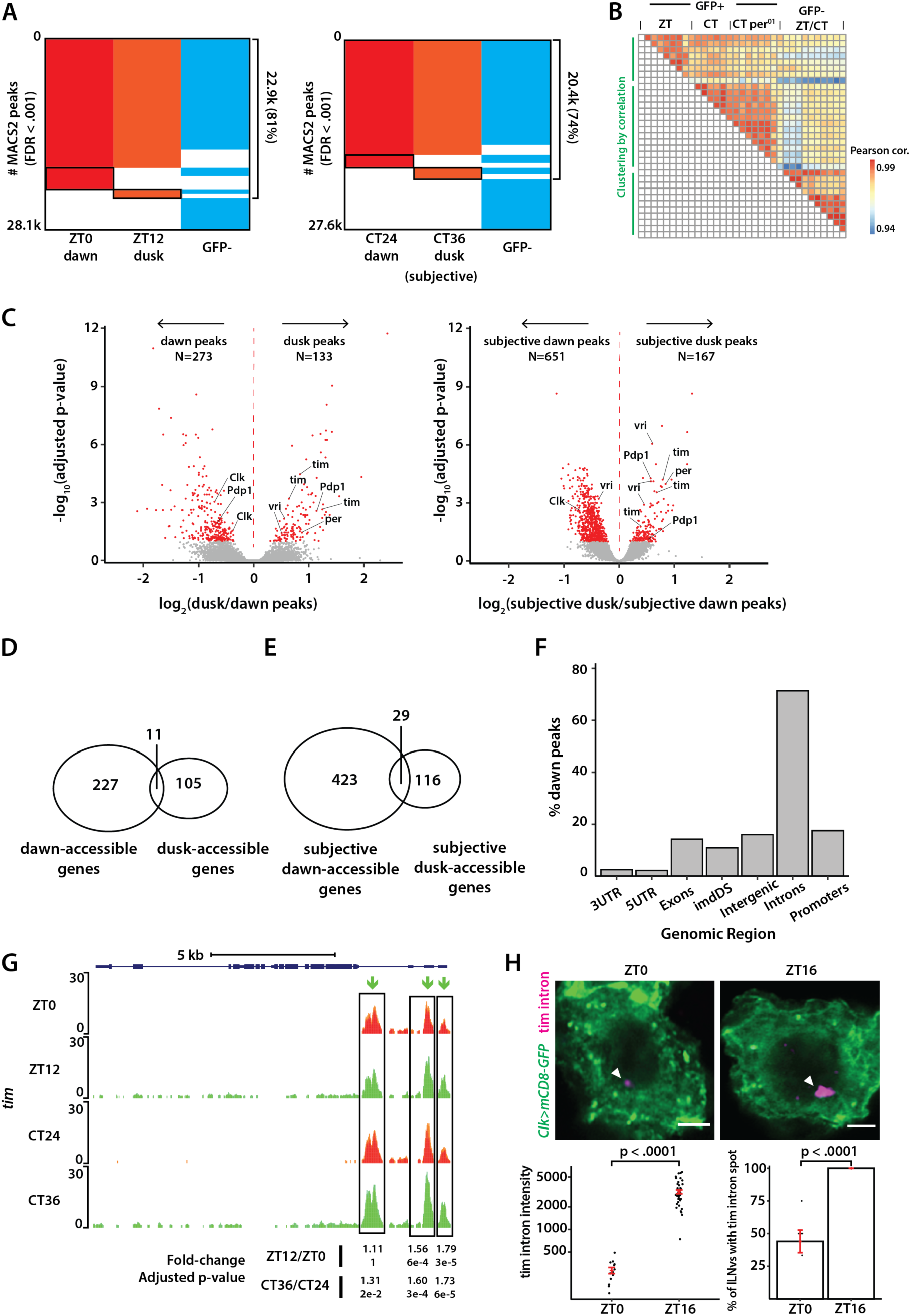
Differential analysis of chromatin accessibility in clock neurons over the circadian cycle. (A) Binary heatmaps of MACS2 called peaks for light-dark cycling (LD: ZT0, ZT12) and constant darkness (DD: CT24, CT36) conditions. MACS2 is used to call distinct ATAC signal peaks for each condition. In total, we identified 28.1k/27.6k peaks under LD and DD conditions. A significant portion of the peaks overlap between GFP-positive clock neurons and GFP-negative cells at both timepoints. Black boxes show peaks called in one time-point only in either LD or DD condition. (B) Pairwise correlation heatmap of all samples used in this study. All sample pairs show high correlation (>0.94). Clock neuron samples in ZT0/ZT12 conditions, in CT24/CT36 conditions (including wildtype and *per^01^*) and non-clock cell samples in both ZT and CT conditions form three distinct clusters. (**C**) Volcano plots of differential peaks under LD and DD conditions. We identified 406 differential peaks under LD and 818 under DD conditions. Peaks corresponding to core clock genes showed higher accessibility at dusk relative to dawn. vri and Pdp1 have more intricate transcriptional regulation as they possess differential peaks at both dawn and dusk. (**D**) Assignment of ATAC peaks to genes by nearest neighbor ranked via chromosome coordinate. The 406 ZT peaks were assigned to 343 distinct genes (∼1.2 peaks/gene) and the 818 CT peaks were assigned to 568 genes (∼1.4 peaks/gene), showing a mostly bijective mapping. (**E**) Distribution of ZT differential peaks with respect to known genomic features by the ChIPpeakAnno Bioconductor package. ‘imdDS’ refers to genomic regions immediately downstream (within 1kb) of promoter regions. (**G**) Normalized ATAC signal pile-up tracks of *timeless* locus under LD and DD conditions. *tim* locus has three regulatory elements near its promoter that are more accessible during dusk when ATAC signal along its full-length gene body could also be identified. (**H**) Representative images of *tim* intron HCR-FISH at dusk and dawn timepoints. Transcription activity of *tim* is higher at dusk consistent with the ATAC differential analysis. Spot fluorescence was quantified and percentage of cells with the transcription spot was counted manually. The statistical test used was a two-sided Student’s t-test. **P < 0.001. Scale bar: 2µm.

To test the validity of our ATAC-seq data, we first assessed whether the chromatin of core clock genes and clock neuron-specific neuropeptide genes exhibited greater accessibility in GFP-positive clock neurons relative to GFP-negative non-clock neurons. Our analysis revealed that the promoters of several core clock genes, including *Clk* and *Tim*, exhibited a higher degree of accessibility in GFP-positive clock neurons as compared to GFP-negative cells (**Figure 1F**). Interestingly, the promoters of other core clock genes, such as *Pdp1* and *Per*, demonstrated accessibility in both GFP-positive clock neurons and GFP-negative cells, suggesting that these promoters are accessible irrespective of their transcriptional status (**Figure S2A**). In these instances, our data revealed that the regulatory intronic regions of these genes exhibited enhanced accessibility in GFP-positive clock neurons relative to GFP-negative cells (**Figure S2A**).

Moreover, we found that the chromatin of several clock neuron-specific neuropeptides, including PDF (expressed in four l-LNvs and four s-LNvs), Trissin (expressed in two LNds), NPF (expressed in two LNds), sNPF (expressed in two LNds and four s-LNvs), AstC-R2 (expressed in a single LNd), ITP (expressed in one LNd and the fifth s-LNv), and DH31 and CNMa (expressed in dorsal clock neurons), displayed enhanced accessibility in GFP-positive clock neurons relative to GFP-negative cells (**Figure 1G**). Notably, many of these neuropeptides are expressed only in a few clock neurons across the whole brain (21), yet our analysis indicates that their gene loci exhibit increased accessibility in clock neurons compared to non-clock cells. These results demonstrate that our ATAC-seq protocol is highly sensitive and can assess genome-wide chromatin accessibility from a relatively small number of FAC-sorted Drosophila clock neurons.

### Chromatin accessibility rhythms over the circadian cycle

To explore changes in chromatin accessibility within clock neurons over the circadian cycle, we analyzed ATAC-seq data from experiments conducted at four distinct time points: dusk (ZT12) and dawn (ZT0) under light-dark (LD) conditions, and subjective dusk (CT36) and subjective dawn (CT24) under constant darkness (DD) conditions. Utilizing the MACS2 peak-calling algorithm (23), we identified over 27,000 peaks from GFP-positive clock neurons across all four timepoints, with a false discovery rate (FDR) threshold of < 0.001 (**Figure 2A**). Under both LD and DD conditions, we observed a significant overlap of over 75% in the peaks between GFP-positive clock neurons and GFP-negative cells, as well as between the two timepoints (**Figure 2A**). This overlap likely corresponds to genes integral to basic cellular functions, encompassing housekeeping genes and those related to general neuronal activities. Only a smaller fraction of peaks is unique to each timepoint under both LD and DD conditions. Nevertheless, our results indicate that GFP-positive clock neurons under LD conditions, GFP-positive clock neurons under DD conditions, and GFP-negative non-clock cells under both LD and DD conditions segregate into three distinct clusters, highlighting distinct chromatin accessibility patterns in each group (**Figure 2B**).

To determine if any of these peaks exhibited differential accessibility across dawn and dusk timepoints, we utilized DESeq2 to quantify changes in chromatin accessibility (24), applying an adjusted p-value cut-off of 0.1 (**Figure S3A**). Of the approximately 27,000 peaks observed under LD and DD conditions, we identified 406 differential peaks under LD conditions and 818 differential peaks under DD conditions (**Figure 2C**). Interestingly, there was only a limited overlap in the differential peaks observed under LD and DD conditions, for instance, between dawn and subjective dawn or between dusk and subjective dusk (**Figures S3B, S3C, S3E**), suggesting that unique chromatin landscapes are being established at each of these timepoints. Furthermore, these observations are consistent with previous RNA-seq findings which indicated that, beyond core clock genes, different sets of genes oscillate under light/dark (LD) and constant darkness (DD) conditions (21, 25).

Out of the 406 differentially accessible peaks under LD conditions, 273 peaks showed higher accessibility at dawn, while 133 peaks showed higher accessibility at dusk. Out of the 818 differentially accessible peaks identified under DD conditions, 651 peaks showed higher accessibility at subjective dawn, while 167 peaks showed higher accessibility at subjective dusk (**Figure 2C**). To understand peak distribution among annotated genes, we assigned peaks to genes by nearest neighbor on 1D-chromosomal coordinates. The 406 differential peaks found under LD conditions were assigned to 343 distinct genes (∼ 1.2 peaks per gene), and the 818 differential peaks found under DD conditions were assigned to 568 genes (∼1.4 peaks per gene) (**Figures 2D, 2E**). Interestingly, we observed that the fold-change difference in peak accessibility is overall higher in LD conditions than in DD conditions (**Figure 2C**). In both LD and DD scenarios, we predominantly found peaks in intronic and promoter regions, consistent with previous reports on the chromosomal localization of regulatory elements (26) (**Figures 2F, S3D**).

Upon examining the chromatin accessibility of core clock genes, we noticed a distinct pattern. Peaks associated with key clock genes like *tim* and *per* exhibited higher accessibility at dusk (ZT12/CT36) (**Figures 2G, S4B**), while peaks associated with the *Clock* gene demonstrated heightened accessibility at dawn (ZT0/CT24) (**Figure S4A**). Specifically, we found that the promoter regions and the intronic regulatory elements of *tim* displayed a higher degree of accessibility at dusk relative to dawn, indicating that these regions transition from an inactive chromatin state to a more active regulatory state over the course of the ∼24-hour circadian cycle. Furthermore, we observed a pervasive opening of chromatin across most of the *tim* gene at dusk timepoints, as indicated by the ATAC signal across its gene body (**Figure 2G**). This could be due to high levels of transcription of *tim* gene in all clock neurons. Conversely, the *per* gene exhibited significant differential peaks primarily in its first intronic region, with increased accessibility at dusk relative to dawn (**Figure S4B**). This intronic region has previously been shown to contain an E-box element which is bound by CLK/CYC proteins (27). We observed a slight increase in the accessibility of the E-box element within the promoter region of the *per* gene at dusk relative to dawn, however, the observed difference did not reach a statistically significant level (**Figure S4B**). Interestingly, we noted that specific genes, such as vri and Pdp1, exhibited regulatory peaks that were highly accessible at both dusk and dawn. (**Figures S4C S4D, S5A**). This pattern may arise from the varied expression of different gene isoforms in specific subsets of clock neurons. For example, from an existing bulk RNA-seq dataset (25), we found that while the short variant of Pdp1 exhibits robust cycling across all clock neuron groups, the long variant is non-rhythmic and is expressed at a consistently high level in a large group of clock neurons (DN1 neurons) throughout the circadian cycle (**Figure S5B**).

Finally, to test if variations in chromatin accessibility indeed correspond to changes in gene transcription, we employed Hybridization Chain Reaction Fluorescence In Situ Hybridization (28) (HCR-FISH) experiments to visualize nascent *tim* mRNA in clock neurons at dawn and dusk. Using a probe set targeting the first intron of *tim*, which is more accessible at dusk relative to dawn, we noticed a more than four-fold increase in fluorescence intensity at dusk versus dawn (**Figure 2H**). Additionally, nearly 100% of clock neurons displayed a *tim* ‘intron spot’ at dusk, compared to lower than 50% at dawn (**Figure 2H**). These findings show a strong correlation between chromatin accessibility and transcription activity for *tim*. Taken together, our study reveals that the chromatin accessibility of regulatory elements of clock-regulated genes oscillates in a circadian manner, with periods of heightened accessibility coinciding with their peak expression times.

### Genes more accessible at dusk contain E-box motifs, while those more accessible at dawn contain VP-box motifs

To identify potential transcription factors that might bind to the differentially accessible peaks within clock neurons, we conducted both de novo and known motif discovery analyses using the HOMER tool (29). Intriguingly, we observed a substantial overrepresentation of the E-box motif (30, 31), with more than 70% prevalence, in peaks that were more accessible at dusk (**Figure 3A**). Conversely, the VP-box motif (15) was predominantly identified in peaks that were more accessible at dawn (**Figure 3A**). The E-box motif is the target of the CLOCK and Arnt-like transcription factors, homologs to the *Drosophila* CLOCK and CYCLE proteins (30, 31), while the VP-box motif can be bound by the transcription factors NFIL3 and DBP, homologs to the *Drosophila* VRILLE and PDP1 proteins (15). Our findings indicate that peaks exhibiting increased accessibility at dusk predominantly correlate with genes regulated by the PER/CLK feedback loop, such as *per*, *tim*, *vri*, and *Pdp1*. Conversely, peaks with enhanced accessibility at dawn tend to be associated with genes regulated by the VRI/PDP1 feedback loop, such as *Clk*. These chromatin accessibility patterns correspond well with known phases of circadian gene expression (21). Next, we compared our ATAC-seq profiles with previously published CLK and RNA polymerase II (Pol II) ChIP-seq data from *Drosophila* clock neurons (32). We observed that differential peak regions for core clock genes either align directly with or are closely adjacent to CLK and Pol II binding sites (**Figure 3B**). This emphasizes the pivotal role these regions play in governing transcriptional dynamics.

**Figure 3.**
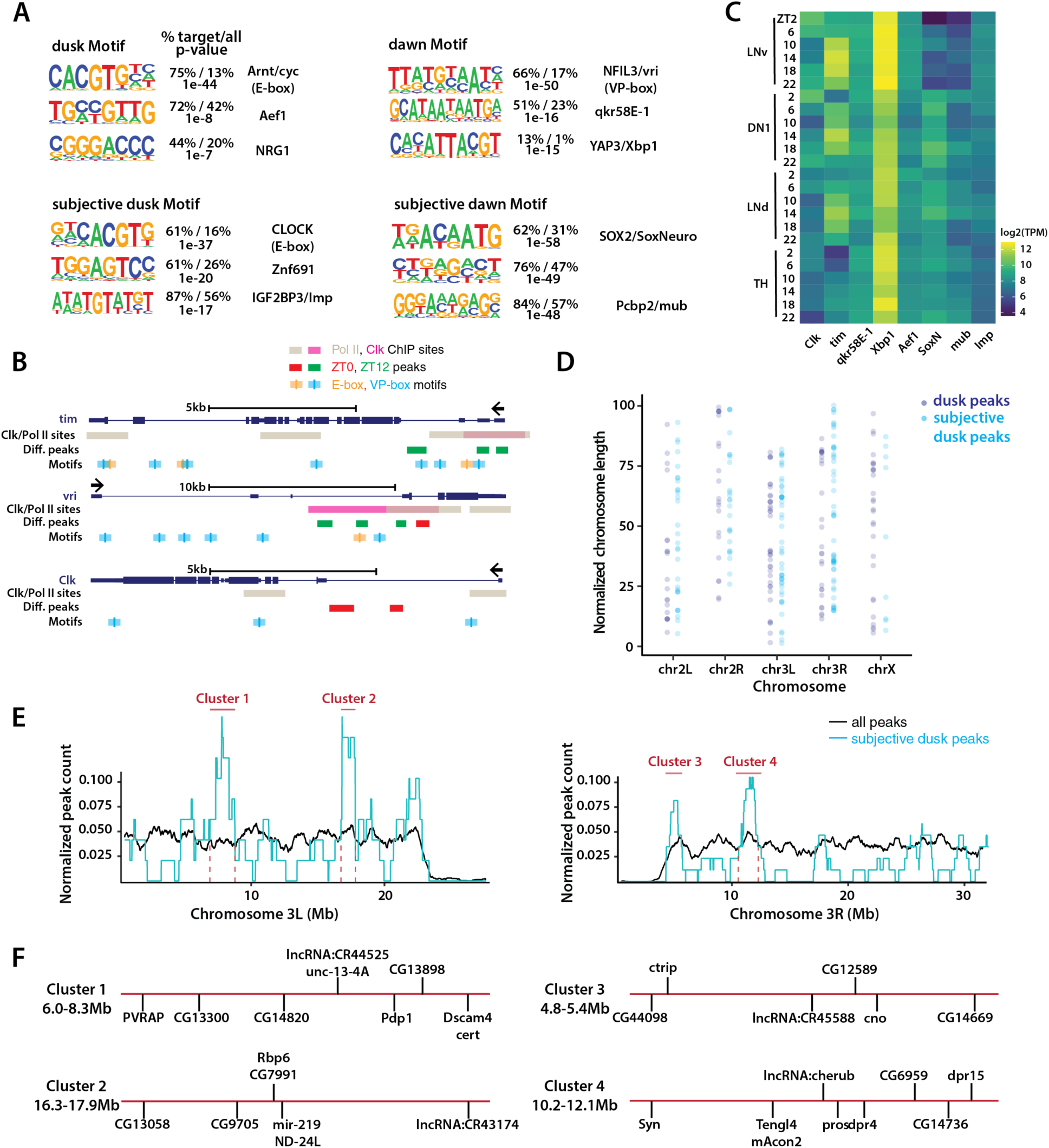
Motifs and clustering analysis of differential peaks and genes. (**A**) De novo motif analysis of differential ATAC peaks. Peaks more accessible at dusk (ZT12/CT36) are highly enriched in E-box sequences, indicating that the corresponding regulatory elements might be directly regulated by the CLK feedback loop. Peaks more accessible at dawn (ZT0/CT24) shows different motif enrichment. ZT0 peaks are enriched in VP-box sequences while CT24 peaks are instead enriched in HMG-domain transcription factor SoxNeuro motif. (**B**) Illustration of E-box and VP-box motif sites along core clock genes *tim*, *vri*, and *Clk*. RNA polymerase II (Pol II) and CLK binding sites (32) and ATAC differential peaks (from this study) are shown. (**C**) mRNA-seq expression profile of identified motif proteins in different clock neuron groups (LNv, DN1, LNd) and a control non-clock neuron group (TH). Analysis was performed with publicly available data (25). (**D**) Distribution of ATAC peaks that were more accessible at dusk (ZT12/CT36) along 1D-chromosome coordinates. Coordinates are normalized by length of the chromosomes (y-axis). (**E**) Zoomed-in distribution of subjective dusk peaks (i.e., peaks more accessible at subjective dusk under DD condition) along chromosomal arms 3L and 3R. Distribution of all identified ATAC peaks is plotted in black showing a mostly uniform distribution. Distribution of subjective dusk peaks is plotted in cyan showing a less uniform landscape suggesting clustering of circadian regulatory elements along 1D-chromosomes. Centromeric regions show near-zero ATAC-signal, as expected due to highly compacted chromatin. Peaks are counted with sliding window bins and count values are normalized to allow comparison between all and differential peaks (see Methods). Clusters which are identified by higher-than-expected normalized peak counts (y-axis) are marked in red. (**F**) Further zoomed-in view of identified gene clusters. Relative positions of the genes are marked with gene symbols on each cluster, normalized by cluster length.

Beyond the E-box and VP-box motifs, we discovered additional motifs linked specifically to peaks that were more accessible either at dusk or dawn. By analyzing a previously published RNA-seq dataset (21), we found that a majority of these transcription factors are expressed across various clock neuron subsets, implying that they could play a role in modulating the diurnal rhythms of essential biological functions (**Figure 3C**). Specifically, at dawn, we found motifs associated with qkr58E-1, a central component of the spliceosome machinery, and Xbp1, a central component in the unfolded protein response (**Figure 3A**). Our findings are particularly striking as past research has demonstrated that alternative splicing of numerous mRNAs crucial for processes like metabolism, cell cycle, apoptosis, and cell proliferation as well as unfolded protein response are regulated rhythmically over the circadian cycle (33, 34). Collectively, our data hint at a coordinated upregulation of vital pathways like alternative splicing and the unfolded protein response during dawn.

Under constant darkness (DD) conditions, we observed a significant enrichment of the E-box motif in peaks that were more accessible at subjective dusk, similar to our previous observations at dusk in LD conditions (**Figure 3A**). However, while in LD conditions we noted a VP-box motif enrichment at dawn, under DD conditions, SOX2 was the predominant motif at subjective dawn (**Figure 3A**). Additionally, we performed known motif analyses, but we couldn’t identify a notable presence of the VP-box motif in peaks with heightened accessibility at this subjective dawn period, suggesting that the VRI/PDP1 feedback loop might be less active or differently regulated under constant darkness conditions. Notably, recent studies have shown that SOX2 is expressed in adult SCN neurons and is essential for the expression of neuropeptides and maintaining ∼24-hour locomotor rhythms (35). Our results underscore the potential pivotal role of SOX2 in modulating circadian rhythms, especially under constant darkness conditions.

Finally, we examined the chromosomal positioning of genes with differential accessibility over the circadian cycle to see if they displayed any specific spatial patterns. Our analysis revealed a distinct spatial pattern across all chromosomes, where genes with higher accessibility at dusk often cluster together and genes that were more accessible at dawn tend to be grouped in proximity along the chromosome length (**Figures 3D, 3E, 3F, S6A, S6B**). Such spatial congregation implies that the physical arrangement of these genes might play a critical role in their co-regulation, thereby driving the ∼24-h circadian rhythms. Collectively, our results reveal that genes with heightened accessibility at dusk are predominantly regulated by the PER/CLK loop through E-box motifs (30, 31), whereas genes with increased accessibility at dawn likely fall under the regulatory domain of the VRI/PDP1 loop, mediated by VP-box motifs (15).

### Chromatin accessibility rhythms correlate with mRNA expression rhythms observed in clock neurons

To assess the relationship between rhythms in chromatin accessibility and gene expression within clock neurons, we compared our ATAC-seq findings with data from a previously published single-cell RNA-seq study (21). Our analysis revealed that between 30-40% of genes displaying rhythms in chromatin accessibility also show cycling patterns in their transcript levels across different clock neuron clusters (**Figure 4A**). Remarkably, despite our ATAC-seq being a bulk procedure that captures all clock neurons, we were able to identify rhythmic chromatin accessibility in genes expressed specifically within smaller subsets of clock neurons. Furthermore, our data highlighted a significant presence of genes linked to the DN1p clock neuron group (**Figure 4B**). This aligns with prior observations indicating a higher prevalence of DN1p neurons in the brain compared to other clock neuron groups (21).

**Figure 4.**
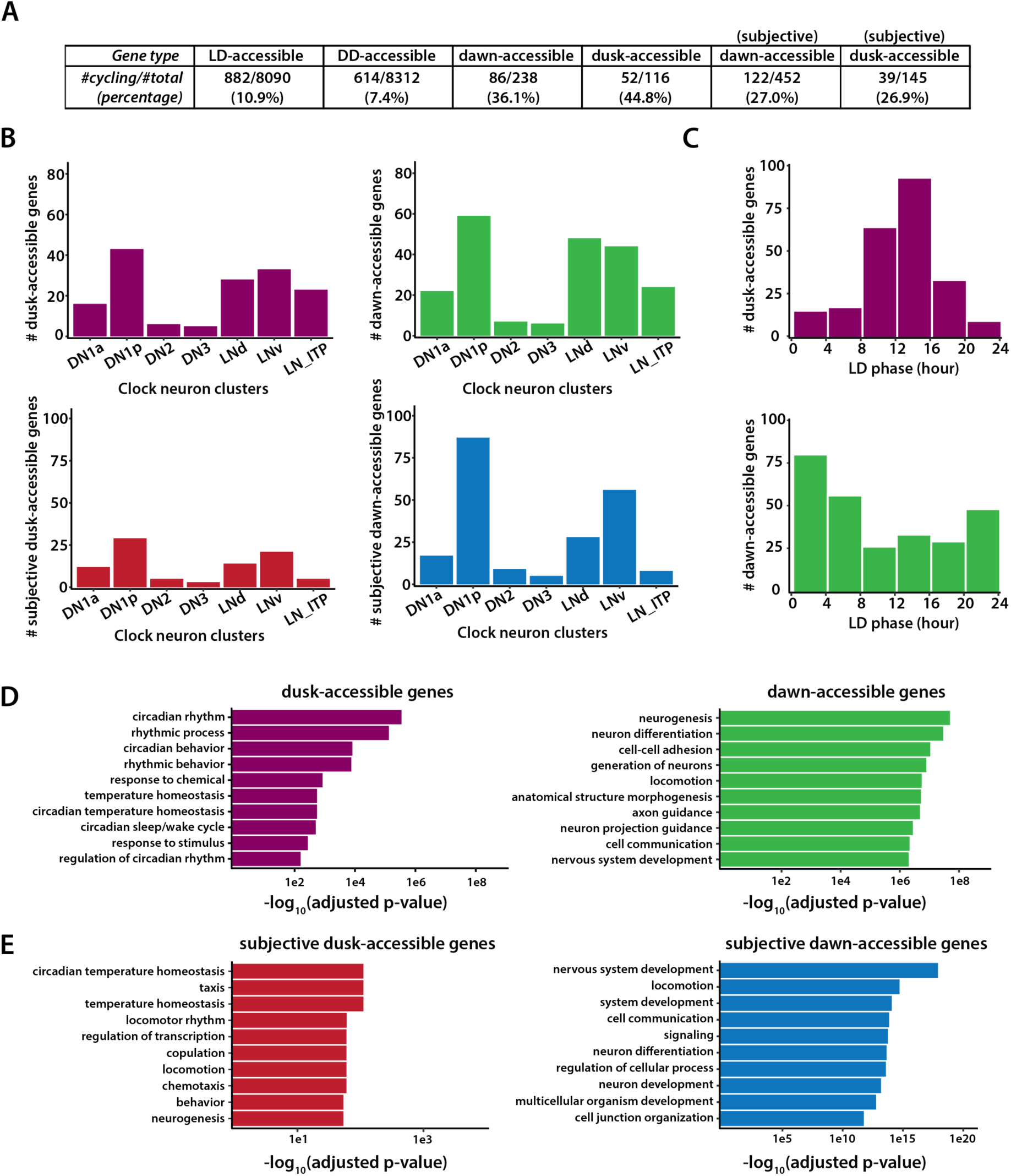
Correlation between gene expression and chromatin accessibility in clock neurons. (A) Table showing overlap between scRNA-seq data set (21) and our ATAC-seq dataset. For genes with any ATAC peak (i.e., genes that are accessible in clock neurons), a small percentage of them are found to be cycling in scRNA-seq dataset (10.9% and 7.4% under LD and DD conditions, respectively). For genes with differential ATAC peaks, overlap percentage increases to 36%/45% under LD and 27% under DD conditions. Analysis was performed with publicly available data (21). (**B**) Distribution of genes with differential ATAC peaks under LD and DD conditions among all the scRNA-seq clock neuron clusters. (**C**) Distribution of phases for peak expression for genes that were more accessible at dawn or dusk. Dawn-accessible genes tend to reach peak transcript levels around ZT0, while dusk-accessible genes tend to reach peak transcript levels around ZT12, indicating strong correlation between chromatin accessibility and transcription. Individual genes may be cycling in multiple clock neuron clusters with different phases and all phases are plotted. (**D**) Ontology enrichment of genes that were more accessible at dusk or dawn. (**E**) Ontology enrichment of genes that were more accessible at subjective dusk or subjective dawn.

Importantly, our ATAC-seq findings reveal a congruence between the phases of higher chromatin accessibility and peak RNA expression as reported in prior RNA-seq studies (25). Specifically, while genes with heightened chromatin accessibility at dusk (ZT12) typically achieve their peak mRNA expression between ZT12 and ZT16, those most accessible at dawn (ZT0) tend to peak between ZT0 and ZT4 (**Figure 4C**). This coherence between enhanced chromatin accessibility and peak gene expression underscores the significance of chromatin dynamics in orchestrating rhythmic gene transcription in clock neurons. The variations in the phases of peak mRNA expression for some genes that we observed could be due to the influence of post-transcriptional regulation (36, 37).

Finally, gene ontology (GO) enrichment analysis revealed that genes with heightened chromatin accessibility at dusk show a significant enrichment for circadian rhythms (**Figures 4D, 4E**). In contrast, genes specifically more accessible at dawn were predominantly associated with processes like neuronal structure morphogenesis and axon guidance (**Figures 4D, 4E**), hinting at an upregulation of these processes in the day’s early hours. These findings are consistent with previous research demonstrating that small ventrolateral neurons (sLNvs), a group of clock neurons, experience daily structural changes leading to more intricate branched projections during the early day compared to the early night (38). To sum up, we found that phases of heightened chromatin accessibility in clock neurons align closely with phases of peak gene expression, underscoring the intricate relationship between chromatin dynamics and rhythmic gene expression within clock neurons.

### Chromatin accessibility rhythms are abolished in the arrhythmic *per^01^* null mutants

We next sought to determine whether a functional clock is required for generating rhythms in chromatin accessibility in clock neurons. To address this, we performed ATAC-seq experiments on FAC-sorted clock neurons from *per^01^;Clk-GAL4;UAS-GFP-NLS* flies (19) at two timepoints under constant darkness, subjective dawn (CT24) and subjective dusk (CT36). Notably, in stark contrast to the wild-type condition where we observed 818 differentially accessible chromatin regions between subjective dawn and subjective dusk, we detected no differentially accessible regions between these timepoints in the *per^01^*mutant flies. Furthermore, the ATAC profiles for core clock genes in the *per^01^*mutants at both timepoints closely resembled the wildtype’s subjective dusk profile, indicating that these genes remain more accessible throughout the circadian cycle in the absence of a functional clock (**Figures 5A, 5D**).

**Figure 5.**
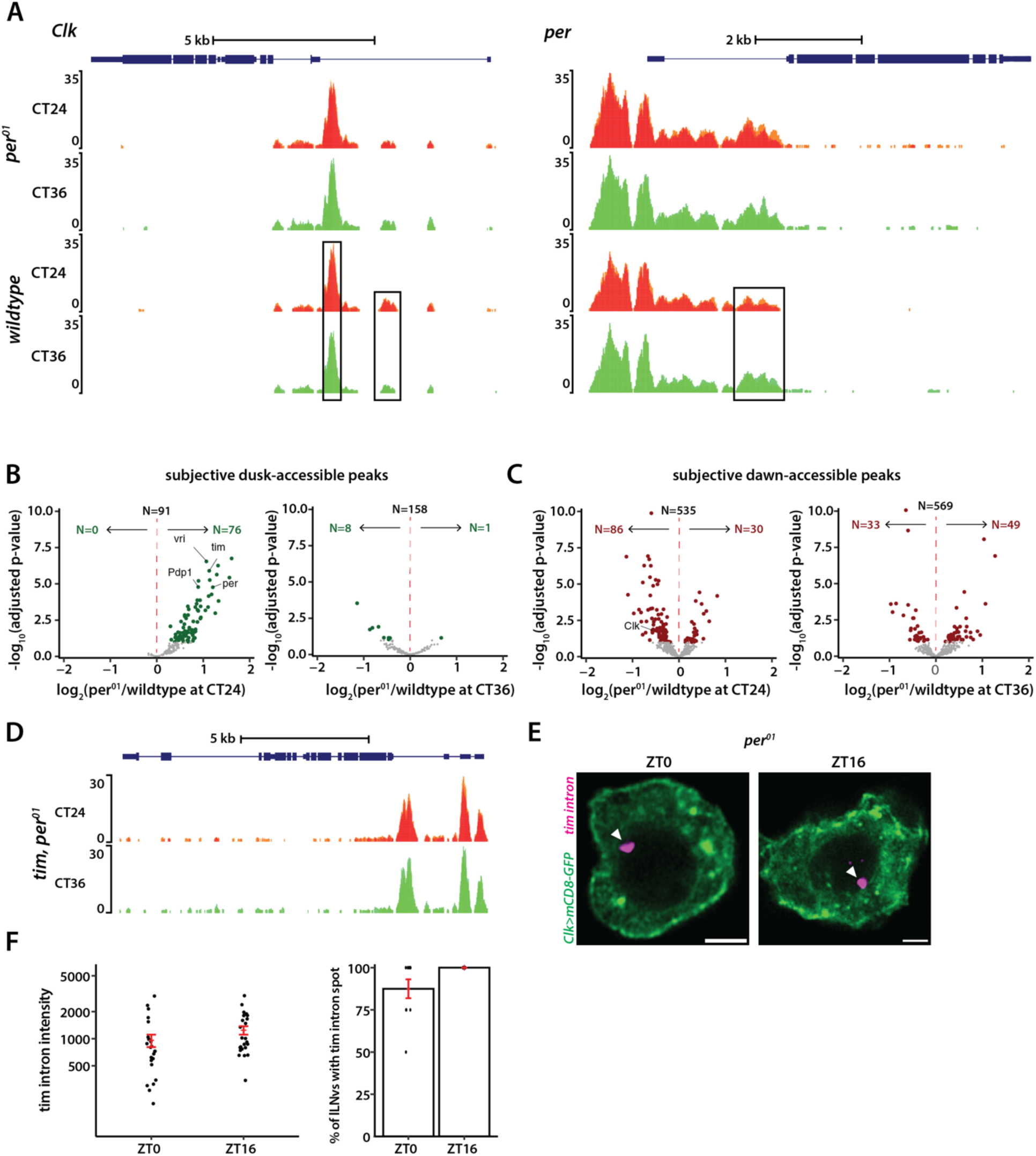
Chromatin accessibility rhythms are abolished in arrhythmic *per^01^* null mutants. (A) ATAC signal pile-up tracks of *Clk* and *per* gene loci in clock neurons from *per^01^* mutants at subjective dawn and dusk. As showcased here and by a genome-wide differential analysis, no differential accessibility is identified between subjective dawn and subjective dusk in *per^01^* mutants. Peaks that are differentially accessible in wildtype are marked by black boxes. (**B-C**) Volcano plots of differential peaks comparing *per^01^* to wildtype at subjective dawn and subjective dusk timepoints. (**D**) ATAC signal pile-up tracks of *tim* locus in clock neurons from *per^01^* mutants at subjective dawn and dusk. (**E-F**) Representative images of *tim* intron HCR-FISH (E) and quantification of fluorescence intensity and percentage of cells with the transcription spot in *per^01^* mutants (F). The statistical test used was a two-sided Student’s t-test, no significance was detected.

Of the 818 peaks with differential accessibility found in wild-type settings, 167 peaks were more open at subjective dusk, whereas 651 were more open at subjective dawn (**Figure 2C**). In *per^01^* mutants, many of these 167 subjective dusk peaks, especially those linked to core clock genes, exhibited increased accessibility at CT24 compared to the wildtype (**Figure 5B**). Notably, almost none showed reduced accessibility in the *per^01^* mutants compared to wildtype at this timepoint. Moreover, when comparing the *per^01^* mutants to wild-type conditions at subjective dusk (CT36), there were no significant differences in chromatin accessibility (**Figure 5B**). Together, our data suggest that the PER protein plays a pivotal role in facilitating dawn chromatin compaction, and clock-regulated genes remain in a consistently “open” state throughout the circadian cycle in the absence of a functional clock.

Next, we examined how the 651 peaks which were more accessible at subjective dawn in wild-type flies are affected in *per^01^* mutants. We observed that peaks associated with the *Clk* gene displayed diminished accessibility at CT24 in the absence of PER, consistent with previous findings of reduced *Clk* mRNA levels in *per^01^* mutants (39) (**Figure 5C**). Yet, a comparative analysis between wildtype and *per^01^* conditions for other peaks did not follow a consistent pattern; some peaks displayed heightened accessibility in *per^01^*, while others were less accessible (**Figure 5C**). Thus, while PER protein plays a specific and significant role in regulating chromatin accessibility of *Clk*, its regulatory control over other dawn-activated genes seems more nuanced, suggesting a possible indirect and context-dependent regulatory mechanism.

As our findings indicate that the lack of PER renders the chromatin of clock-regulated genes more accessible, we asked if this leads to higher transcription across the circadian cycle. To test this, we employed HCR-FISH to visualize nascent *tim* mRNA transcripts in *per^01^*background. Unlike the wild-type scenario (**Figure 2H**), we observed no significant difference in the fluorescence intensity of the *tim* ‘intron spot’ between dusk and dawn timepoints in *per^01^* mutant flies (**Figures 5E, 5F**). Moreover, nearly all clock neurons in *per^01^* mutant flies exhibited a *tim* ‘intron spot’ at both timepoints over the circadian cycle (**Figures 5E, 5F**), suggesting that *tim* is expressed at high levels throughout the day in the absence of a functional clock. To sum up, our results indicate that a functional clock is essential for the chromatin compaction of clock-regulated genes during the repression phase, which is required for generating ∼24-hour gene expression rhythms.

### Differential chromatin accessibility in non-clock neurons

We asked whether we could identify any differentially accessible peaks between dawn and dusk timepoints in GFP-negative cells, which include a diverse set of neurons and glial cells in the brain. Interestingly, we detected a small group of differentially accessible peaks that were more accessible at dusk (ZT12), a period typically associated with sleep in flies, in comparison to dawn (ZT0) (**Figures 6A, 6B**). This suggests that these regulatory elements might be more accessible across a broad range of neurons in the brain during dusk. These peaks are associated with a select group of genes, incuding the eukaryotic translation initiation factor (*eIF5B*), Ecdysone-induced protein 75B (*Eip75B*), Shaker (*Sh*), Vrille (*vri*), and *CG31140* (**Figures 6A, 6B**). Given the crucial role of eIF5B in translation initiation, our results suggest a potential upregulation in translation activity during dusk, which aligns with previous studies that showed an accumulation of messenger RNAs and proteins related to translation during the sleep phase (40). The vrille gene, required for both development (41) and circadian rhythms (42), encodes a transcriptional repressor and is known to be widely expressed in the brain. We found that ATAC-peaks linked to the vrille gene exhibited heightened accessibility at dusk not just within clock neurons but also in other brain neurons (**Figure S4D**). We also observed increased accessibility at dusk in sites connected to CG31140, a lipid kinase essential for the ATP-driven phosphorylation of diacylglycerol (DAG) to yield phosphatidic acid (43), hinting at temporal regulation of lipid signaling pathways.

**Figure 6.**
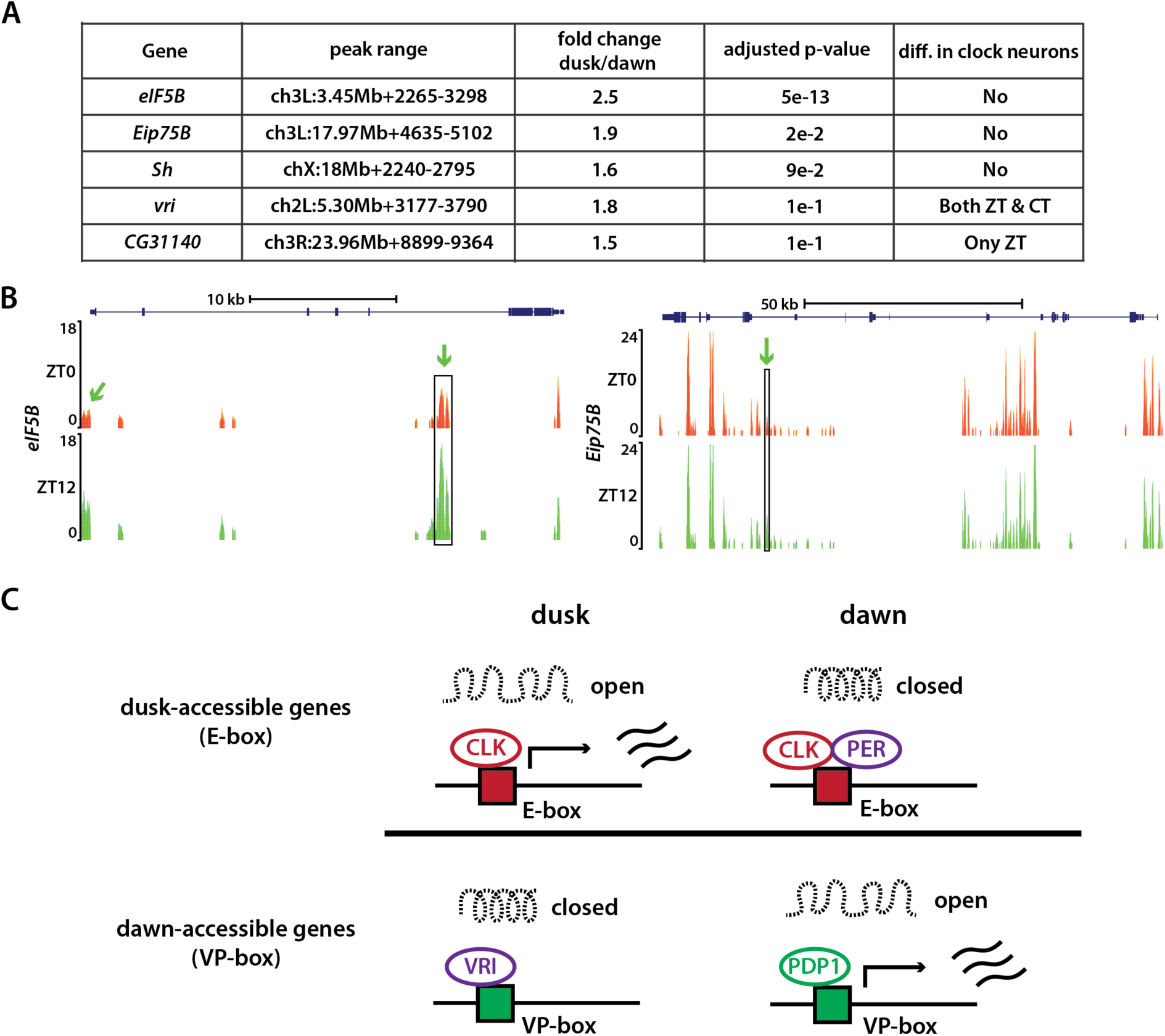
Differential chromatin accessibility in non-clock cells and model. (A) Summary table of differential peaks identified in non-clock cells at dusk compared to dawn. (B) ATAC signal pile-up tracks of *eIF5B* and *Eip75B* gene loci in non-clock cells at dawn and dusk. Differentially accessible peaks are marked in black boxes. (**C**) Summary of our model. Chromatin accessibility of regulatory elements of clock-regulated genes oscillates within a 24-hour day. E-box containing genes were typically more accessible at dusk, while VP-box containing genes were typically more accessible at dawn.

Interestingly, our analysis revealed enhanced accessibility in the regulatory regions of two key sleep-regulating genes, Shaker and Eip75B (44–46), during dusk—a period when flies are naturally asleep—compared to dawn. The Shaker gene, responsible for encoding a voltage-gated potassium channel (47), plays a critical role in modulating the excitability of neurons and mutations in Shaker have been linked to disrupted sleep behaviors, including shorter sleep and more fragmented sleep patterns (44, 45). Recent studies have shown that the ecdysone receptor (EcR) and its downstream nuclear hormone receptor Eip75B (E75) influence the timing and duration of sleep (46). The heightened accessibility of regulatory elements associated with Shaker and Eip75B at dusk offers tantalizing hints towards the intersection of chromatin dynamics and the molecular underpinnings of sleep regulation.

## Discussion

Biological clocks are regulated by clock proteins that participate in complex feedback loops, which ensure the timely expression and repression of specific genes that aligns with the day-night cycle. Our study introduces an additional layer of complexity to this understanding. We have unveiled how clock proteins coordinate rhythms in chromatin accessibility across the regulatory elements of numerous clock-regulated genes in *Drosophila* clock neurons, which might be vital for the establishment of approximately 24-hour rhythms in mRNA expression (**Figure 6C**). Our results suggest that differences in chromatin accessibility could be a key factor that contributes to the observed heterogeneity in circadian gene expression within clock neurons. A recent study on mouse liver cells has demonstrated similar dynamic changes in the accessibility of the liver genome throughout the day (48). This convergence of findings—both in Drosophila and mammals—underlines the universality and importance of chromatin accessibility in shaping circadian rhythms.

### Chromatin accessibility rhythms in clock neurons and non-clock cells

Our study identified two distinct gene sets based on their chromatin accessibility patterns: one set has regulatory elements that were more accessible at dusk, while the other set has regulatory elements more accessible at dawn. Motif analysis revealed that genes with heightened accessibility at dusk possessed E-box motifs (30, 31), suggesting they might be governed by the PER/CLK loop. In contrast, genes more accessible at dawn possessed VP-box motifs (15), suggesting they might be controlled by the VRI/PDP1 loop. In addition to the E-box and VP-box motifs, we identified other motifs uniquely associated with chromatin peaks exhibiting heightened accessibility at either dusk or dawn. Intriguingly, peaks with heightened accessibility at dawn were associated with motifs tied to alternative mRNA splicing and the unfolded protein response. These findings align with prior studies and emphasize the cyclical regulation of processes like alternative splicing and the unfolded protein response over the circadian cycle (33, 34).

In our analysis of chromatin accessibility in GFP-negative (non-clock) cells, we made an intriguing discovery that certain gene-associated peaks exhibited increased accessibility at dusk compared to dawn. This suggests a unique temporal regulation in the chromatin landscape of these specific genes across a broad range of neurons in the brain during the day-night cycle. It’s worth noting that the opposite scenario – peaks being more accessible at dawn than at dusk in non-clock cells – was not observed. Considering that flies are diurnal, they sleep during the lights-off phase from ZT12 to ZT0, it would be intriguing to explore whether these chromatin accessibility changes play a role in regulating sleep patterns, behaviors, or other physiological processes during their sleep phase. Future investigations in this direction could provide intriguing insights into the potential regulatory mechanisms at play during the sleep phase of these organisms. Furthermore, it’s intriguing to consider how these rhythms emerge without a traditional clock mechanism and to discern the influence of ambient light, or its absence, on these chromatin accessibility patterns.

### Chromatin accessibility correlates with peak mRNA expression in clock neurons

The relationship between chromatin structure and gene expression can be intricate and multifaceted, as various factors, such as the timed presence of specific transcription factors during the circadian cycle, epigenetic alterations, and the spatial organization of the nucleus, might all modulate gene expression. Interestingly, our research demonstrates that a significant 30-40% of genes displaying circadian rhythms in chromatin accessibility also show oscillations in transcript levels among various clock neuron clusters. This underscores the crucial interplay between chromatin dynamics and gene transcription in the case of clock-regulated genes in clock neurons. This is especially striking considering the central role post-transcriptional regulation plays in shaping ∼24-hour circadian rhythms (36, 37).

Circadian regulation has been shown to be largely cell- and tissue-specific in both *Drosophila* and mammalian clock neurons (21, 49). In most cases, only transcripts of core clock genes and a handful of other genes demonstrate consistent cycling across all clock neuron groups. While many mRNAs showed cyclical patterns in single-cell RNA-sequencing studies (21), these patterns were often absent in the wider context of fly head or brain mRNA (50). However, our ATAC-seq method proved effective in detecting rhythmic chromatin accessibility in gene regulatory regions, even if rhythmic expression was limited to a small subset of clock neurons. Finally, our observation that genes with similar accessibility patterns tend to cluster along the chromosomal arms might suggest coordinated chromatin remodeling or a higher-order chromosomal organization that facilitates or restricts access to certain genes at specific times. Future studies can delve deeper into the implications of such spatial arrangement on the functionality of the circadian rhythm and the underlying processes.

### How might PER protein drive chromatin compaction during the repression phase?

Past studies have shown that the activators, CLOCK and BMAL1, interact with histone acetlytransferases (HATs) such as p300 and CBP (51), as well as the methyltransferase MLL1 (52), to promote acetylation and methylation of histones, respectively, to promote an open chromatin state to facilitate transcription. Intriguingly, CLOCK has also been reported to have intrinsic HAT activity (53). On the other hand, PER/CRY complex is known to be a large macromolecular complex composed of at least 25 proteins (54, 55). PER protein has been shown to recruit several transcriptional repressor complexes, such as Mi-2/nucleosome remodelling and deacetylase (NuRD) (56), SIN3-HDAC (57), and Hp1γ–Suv39h histone methyltransferase (58), as well as RNA helicases DDX5 and DHX9 (59), to target genes to promote repression. The interaction of PER with these transcriptional repressor complexes may promote chromatin compaction, essential for the effective silencing of target genes and the creation of rhythmic gene expression patterns vital to circadian regulation. Subsequent studies could delve deeper into the molecular mechanisms controlling chromatin accessibility rhythms within clock neurons.

Chromatin accessibility plays a critical role in regulating gene expression and determining cell fate. As animals develop, distinct DNA regions become variably accessible to transcriptional machinery, ensuring timely and appropriate gene expression at the right stages of development (3, 4). Although chromatin configuration, typically set during development, tends to remain stable throughout the lifespan of the cell (3, 4), our research presents divergent findings in clock neurons. Our research uncovers a dynamic chromatin landscape in clock neurons: chromatin accessibility of regulatory elements of clock-regulated genes is not static but rather oscillates within a 24-hour day, is dependent on a functional clock, and this rhythmic pattern recurs daily.

## Materials and Methods

### Fly stocks

Flies were raised on standard cornmeal/yeast media and maintained at room temperature (20 to 22 °C) under a 12h:12h LD schedule. The following flies used in the study were previously described or obtained from the Bloomington Stock Center: *Clk-GAL4* (*20*)*, per01* (*19*). Flies were entrained to Light-Dark (LD) cycles where they were exposed to 12-hour Light-Dark (LD) cycles for 5-7 days, followed by a shift to complete darkness (DD) for another 6-7 days. The initiation of the light phase is labeled as Zeitgeber Time (ZT) 0, while ZT12 signifies the beginning of the dark period. When referencing the times in the continuous darkness phase, we use Circadian Time (CT) - with CT0 (subjective dawn) marking the time when lights would have been turned on and CT12 (subjective dusk) the time when lights would have been turned off. The labels DD1 and DD2 represent the first and second days of complete darkness, respectively. To make the UAS-GFP-NLS flylines, coding sequences of GFP-NLS-tetR fusion protein was synthesized and cloned into UAS expression plasmid pJFRC-MUH (Addgene #26213) by GenScript Biotech, and this plasmid was microinjected into phiC31 integrase line with docking site attP40.

### Hybridization Chain Reaction Fluorescence In Situ Hybridization (HCR-FISH)

To perform Hybridization Chain Reaction Fluorescence In Situ Hybridization (HCR-FISH) in whole-mount Drosophila brains, we adapted our previous protocol on single molecule RNA-FISH (60). Probes and amplifier hairpins were synthesized by Molecular Instruments. For our HCR-FISH experiments, we housed the flies in density-regulated food vials, each containing 4 females and 4 males. They were acclimated in incubators for 5-7 days. All imaging tests were conducted on male or female flies aged between 5-7 days. Notably, there were no observable differences in the outcomes of our experiments between genders. Specific fly genotypes and Zeitgeber Time (ZT) details are provided in the accompanying figure legends. To express transgenes in the brain’s clock neurons, we employed the GAL4/UAS system. We captured our images using the Zeiss LSM800 confocal microscope, equipped with an AiryScan super-resolution module that offers 125 nm lateral and 350 nm axial resolution. Imaging was done utilizing a 63x Plan-Apochromat Oil (N.A. 1.4) objective, with laser lines of 405, 488, and 561 nm. Our imaging sessions included collecting Z-stack series (with approximately 250 nm per Z-slice) of distinct clock neurons. Following image acquisition, we imported the CZI files into ImageJ, ensuring we maintained a lossless 16-bit resolution for each channel, using the Bio-Formats Importer to process them as composite images. Subsequently, we manually delineated regions of interest (ROIs) on the channel, fine-tuning the white values to optimize visualization. To guarantee an unbiased fluorescence analysis, we remained consistent with our visualization settings across all images, keeping them specific to each fly line and neuron type. To measure fluorescence intensity of HCR-FISH spot, we used the ImageJ software to measure both the mean pixel brightness (in arbitrary units or a.u.) and the geometric area (in µm^2) of the designated ROI. To calculate the integrated intensity of the spot, we multiplied the pixel brightness with the geometric area, resulting in a numeric value represented in a.u.×µm^2.

### Brain dissociation and flow cytometry

To obtain suspension of clock neurons, we adapted steps described in (22). Flies were anesthetized and brains were dissected for up to an hour with Schneider’s medium (Sigma S0146) supplemented with 1% bovine serum albumin (BSA, Sigma A7030). Brains were kept on ice, and we aimed to get 60 brains per sample. After dissection, we added collagenase (Sigma C0130) to a final concentration of 2mg/mL. Samples were incubated at 37C for 20min without mixing. We then adjusted solution volume to 200uL and triturated by pipetting with standard 200uL tips 1Hz for up to 80-100 times. Cells were resuspended in 300uL PBS with 0.1% BSA by centrifugation at 300g for 5min at 4C. To obtain a positive control for dead cell marker, suspension equivalent to 3 brains were heated at 60C for 5min. DAPI was added at a final concentration of 2ug/mL as dead cell marker. Finally, suspensions were filtered using FACS tube with strainer cap (Falcon 352235) to obtain single cell suspension and live clock neurons were sorted with FACS sorter (BD Aria III). Cells were sorted into 1.5mL DNA low-bind tubes (Eppendorf) with 50uL PBS with 0.1% BSA. DAPI threshold was set based on the dead cell control sample and GFP threshold was set based on the expected fraction of clock neurons in the whole brain (0.1%). We typically obtained 1500-3000 clock neurons per sample. For non-clock cell control samples, we sorted ten thousand live cells of similar sizes. BSA solution was filtered with 0.22um polyethersulfone membrane (Millipore GPWP047000) and solutions were kept at 4C and reused for up to 5 days to prevent potential contamination. All plastics (including tubes and tips) were coated with PBS 1% BSA for all steps starting trituration to prevent cell loss.

### Assay for transposase-accessible chromatin by sequencing (ATAC-seq)

We adapted steps described in (18, 22). Below, we describe our experiment and analysis procedure.

#### Transposition reaction

Sorted clock neurons were immediately used for ATAC assay after sorting. All plastics were pretreated with 1XPBS with 1% BSA to prevent cell/nucleus loss. First, cells were spun down at 600g for 8min, 4C using spin-out rotor centrifuge. Supernatant was carefully pipetted out and cells were resuspended in 50uL of ice-cold nuclei lysis buffer (10mM Tris-Cl pH 7.4, 3mM MgCl2, 10mM NaCl, 0.1% IGEPAL CA-630) by gently pipetting a few times on ice. Next, nuclei were pelleted at 1200g for 12min, 4 °C. Supernatant was carefully pipetted out and permeabilized nuclei were resuspended in 10uL of transposition solution (5uL 2xTD buffer, 0.5uL loaded Tn5 transposase, 4.5uL ultrapure water) by gently pipetting. 2xTD buffer and Tn5 transposase were commercially available (Diagenode C01019043, C01070012). Reactions were incubated at 37C for 30min without agitation, after which DNA was purified by silica column (Zymo DCC-5) with final volume of 20uL using elution buffer. Elution buffer was heated to 55 °C and we performed two elution steps (10uL each) to increase yield. Purified DNA was stored at -20 °C for up to one week before library preparation.

#### Library preparation and sequencing

Transposed DNA was adjusted to volume of 20uL for each sample and limited-cycle PCR mix was prepared (20uL transposed DNA, 5uL single-indexed Nextera primer mix at 10uM each, 25uL NEBNext High-fidelity PCR master mix). We purchased Nextera primer mix from Integrated DNA technologies which provides universal 8bp index sequences. Next, we performed pre-amplification to repair nicks and then determined amplification cycle numbers by qPCR following standard procedure. We made sure that amplification is in exponential phase and typically 8-10 additional cycles were used to obtain sufficient product for sequencing (>5nM in 20uL after purification). Amplified libraries were purified by non-selective SPRI bead purification and size distribution was assessed by gel electrophoresis (Agilent Tapestation, high-sensitivity DNA 5000 assay). If excess adapter dimers were observed, an extra size-selective SPRI purification was performed. Libraries were then sequenced by shared NovaSeq S4 flow cell, paired-end 150bp cycle targeting 30-50 million reads per sample.

#### Alignment and quality controls

Reads were trimmed to remove sequencing adapters (cutadapt 4.1) and aligned against the dm6 genome assembly using Bowtie2 (version 2.4.5) with parameters ‘--very-sensitive --dovetail -X 1000’. After alignment, mitochondrial reads were removed (samtools 1.15.1) and PCR duplicates were removed using Picard Tools (version 2.27.4). MultiQC (version 1.13a) was used to aggregate result summaries and ATACseqQC (version 1.21.0) was used for standardized quality control designed for ATAC-seq. We typically observed an alignment rate of 30-50%. Unaligned fragments show contamination enriched in bacterial and fungal sequences (NCBI BLAST). Mitochondrial DNA (mtDNA) reads were around 3-5% and PCR duplicate rate was typically around 50%. Overall, we obtained 6-15 million unique non-mtDNA aligned reads.

#### Peak calling and differential analysis

Alignment BAMs were converted to BED with Rsamtools (version 2.13.0) and MACS2 (version 2.2.7.1) was used for peak calling with typical ATAC analysis parameters ‘-f BED -g dm -q 0.001 –nomodel --shift -37 --extsize 74 --min-length 74 --max-gap 74 --keep-dup all’. To quantify chromatin accessibility of each peak, we counted read ends that map within the peak using a publicly available R custom package edgeCounter (https://github.com/yeyuan98/edgeCounter). Counts were then used for differential analysis with DESeq2 (version 1.38.3). Pile-up tracks are publicly available on UCSC genome browser (https://tinyurl.com/2ftuhv5p).

### Additional bioinformatics analyses

HOMER (version 4.11) was used for de-novo and known motif enrichment analysis. gProfiler was used for gene ontology enrichment in genes containing differential peaks. All genes showing ATAC signal were used to scope the statistical domain to remove non-specific enrichment. HOMER de novo analysis results are publicly browsable (https://yeyuan98.github.io/motifAnalysisExports/). Pairwise correlation was computed by counting ATAC reads in 10kb genomic bins. Other analyses shown in this study were performed with packages from the R Bioconductor project (61) including genomic feature annotation using ChIPpeakAnno (62). Published RNA-seq data of LNv, DN, and LNd (25) was aligned against the FlyBase Release 6.45 assembly using STAR aligner. List of identified rhythmic genes in each cluster from a published single cell RNA-seq data (21) was used to compare our ATAC-seq results to scRNA-seq data.

### Analysis of locomotor activity and rhythmicity

We investigated the locomotor activity of individual adult male flies (3-5 days old) using TriKinetics DAM2 Drosophila Activity Monitors. Each fly was placed in a glass capillary tube (around 4 mm in diameter and 5 cm long) containing food made of 2% agar and 4% sucrose. These monitors, equipped with infrared sensors, detect when the flies move across the tube’s midpoint, causing infrared beam breaks. The monitors were stationed in incubators, and as the flies moved, their activity, measured as infrared beam breaks, was recorded on a connected computer. For behavior analysis, we divided this activity data into 30-minute intervals. To process and visualize this data, we utilized both ClockLab (Actimetrics) software. Each fly’s activity was normalized such that its average daily activity (over 48 intervals) equaled 1. By computing the population mean of this normalized activity, we produced normalized activity plots which are showcased in the figures. For analyzing rhythmicity, we assessed the activity counts from each fly during the total darkness phase post-entrainment. Using the ClockLab software, we employed a chi-square periodogram analysis (confidence level set at 0.001) to determine the free-running period of the circadian clock and each fly’s rhythmicity. The results from this analysis, the “Power” and “Significance” metrics, enabled us to derive a “Rhythmic Power” value, representing the robustness of each fly’s rhythm.

### Statistical analysis

For assessing the characteristics of HCR-FISH spots, including fluorescence intensity and the percentage of cells with spots, data were aggregated from multiple hemi-brain images. These images were taken across over three separate experiments, ensuring a robust pool of biological replicates. When measuring fluorescence, each neuron was unique to a brain and was measured only once. For behavioral studies, the sample size ranged between 30 to 60 flies, only makes flies were used for behavior experiments. For ATAC-seq experiments, we dissected ∼60 brains from both male and female flies. We repeated each experiment four times. All data was plotted and statistically analyzed using OriginPro from OriginLab (Northampton, MA, USA).

## Acknowledgments

We thank the Bloomington Drosophila Stock Center for providing fly strains. We acknowledge support from the University of Michigan Biomedical Research Core Facilities (Flow Cytometry Core and Advanced Genomics Core). We thank Yangbo Xiao for generating NLS-TetR-GFP flies and Rafael De Gouvea for help with preparation of figures and members of Yadlapalli lab for help with brain dissections and for discussion and comments on the manuscript. SY is supported by the National Institutes of Health R35 grant (R35GM133737). SY is an Alfred. P. Sloan Fellow, and McKnight Scholar. MVB was supported by the NIH Cellular and Molecular Biology Training Grant (T32-GM007315) and an F31 from NINDS (F31-NS127484). EJC is a McKnight Scholar, Rita Allen Milton Cassell Scholar, and Pew Biomedical Scholar.

## Author Contributions

YY and SY conceived and designed the study. YY performed all the ATAC-seq experiments and analyzed data in consultation with MB, JC, and SY. QC performed HCR-FISH experiments and confocal imaging. SY supervised the project and data analysis. YY and SY wrote the manuscript with input from all authors.

---

**Supplementary Figure S1.**
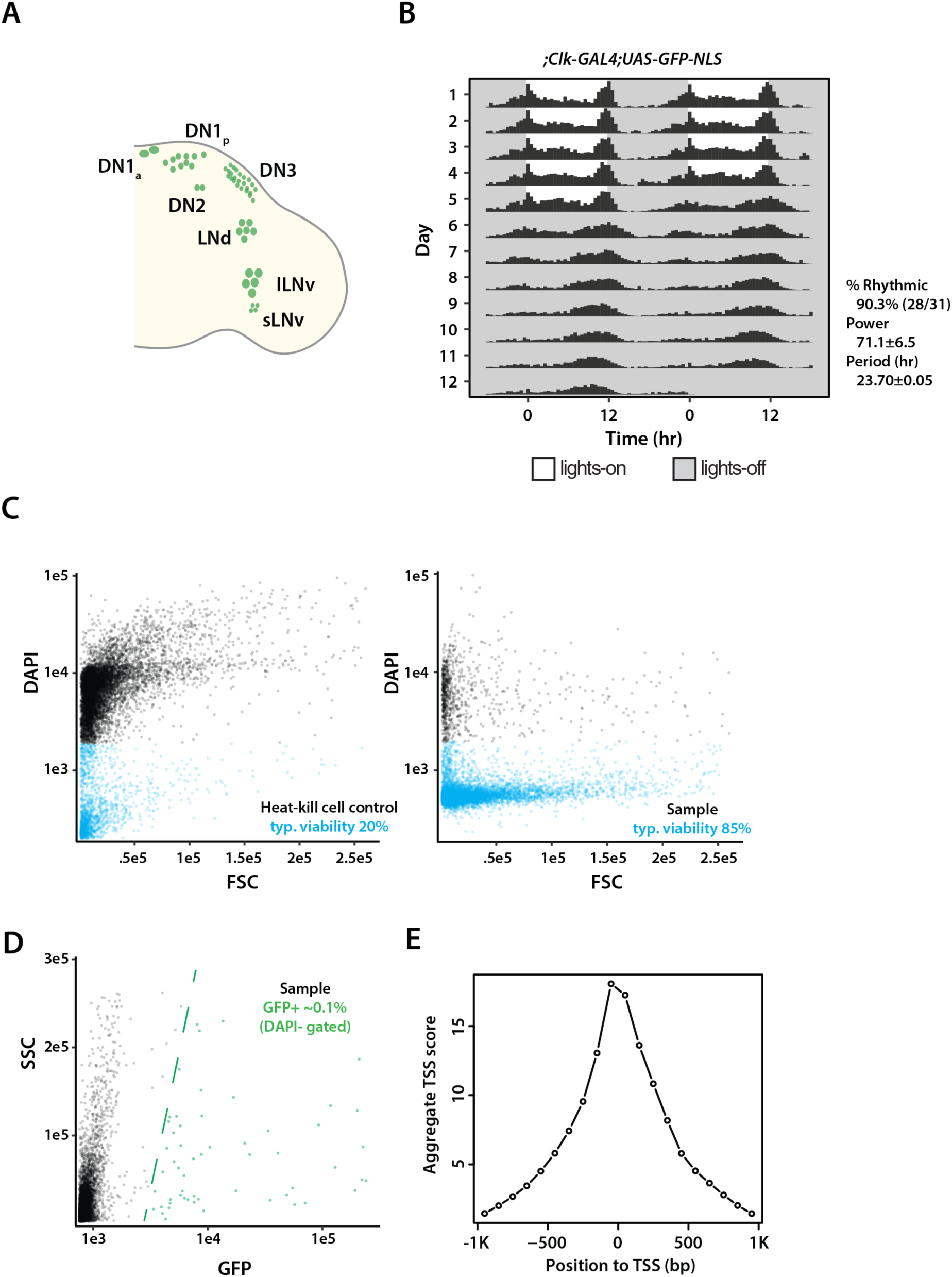
Experimental schema for entrainment and FACS procedure. (**A**) Schema of *Drosophila* clock neuron subgroups shown in a hemi-brain. (**B**) Behavior actogram of *Clk-GAL4>UAS-GFP-NLS* flies. These flies were entrained to LD cycles (ZT0: lights on; ZT12: lights off) for 5 d and released into DD for 7 d. Averaged population locomotor-activity profiles of flies (n = 31) in LD and DD with rest-activity shown for two consecutive days in the same line. These flies display rhythmic behaviors with a period of 23.70 ± 0.05 h, with activity peaks around the time of lights on and lights off. (**C**) Representative plots for the fluorescence activated cell sorting procedure. Left panel shows DAPI-forward scatter (FSC) plot for a typical dead-cell control. Right panel shows a representative DAPI-FSC plot for experiment samples. Typical cell viability is 80∼90%. (**D**) Representative side scatter (SSC)-GFP plot where live (DAPI-) cells are gated by GFP fluorescence. GFP threshold for clock neurons is set based on ∼0.1% expected positive rate. Clock neurons show significantly higher GFP signal, typically >10-fold compared to the GFP-negative population. Green line shows a representative GFP-gate setting. (**E**) Representative TSS enrichment score distribution showing clear signal enrichment immediately upstream of TSS. Enrichment score is computed using the ATACseqQC package.

**Supplementary Figure S2.**
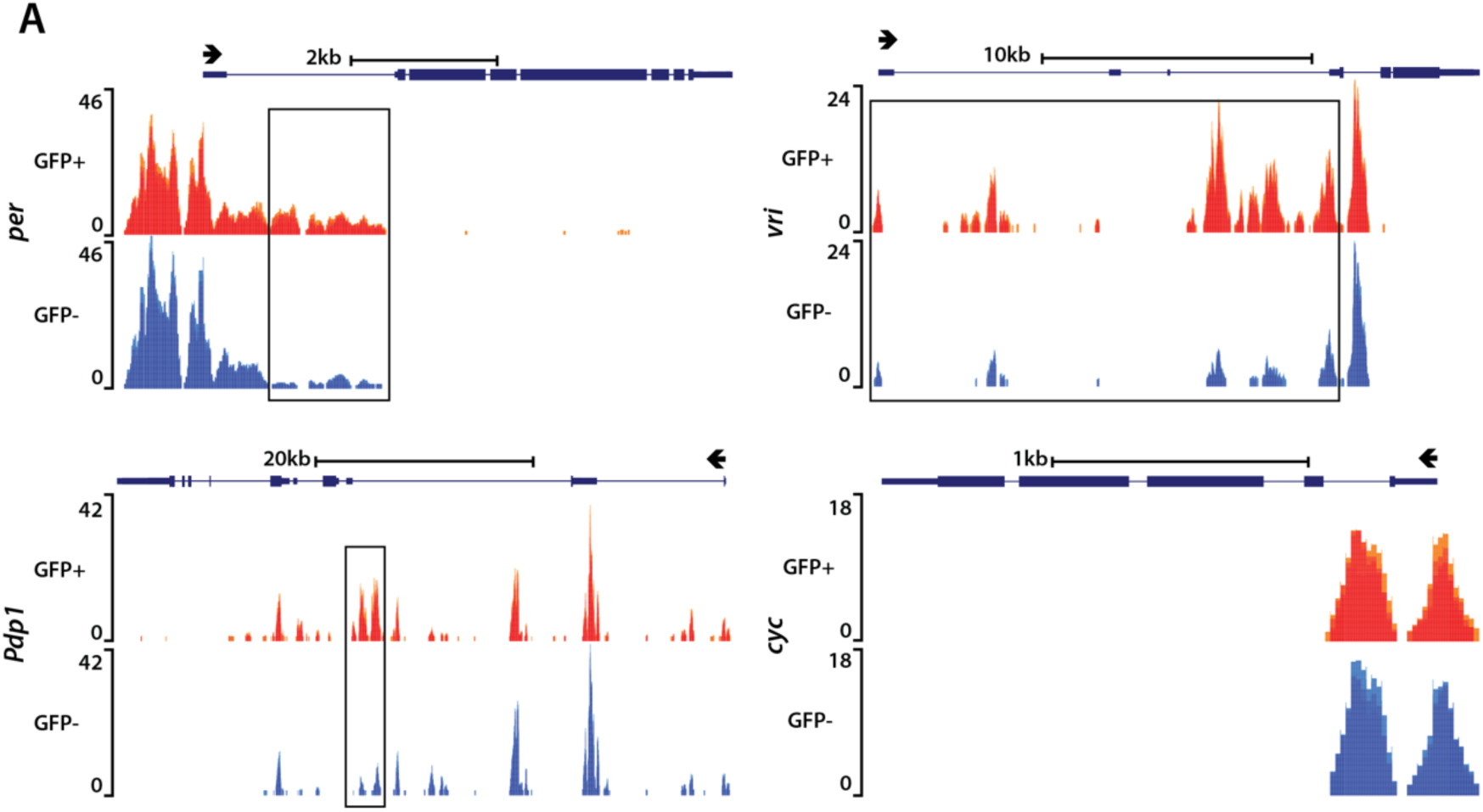
ATAC signal pile-up tracks for core clock genes in GFP-positive clock neurons and GFP-negative non-clock cells. (A) ATAC signal pile-up tracks at *per, vri, Pdp1* and *cyc* loci in clock neurons (GFP-positive) and non-clock cells (GFP-negative). We did not observe any accessibility changes in the *cycle* locus. Differentially accessible peaks are marked in black boxes.

**Supplementary Figure S3.**
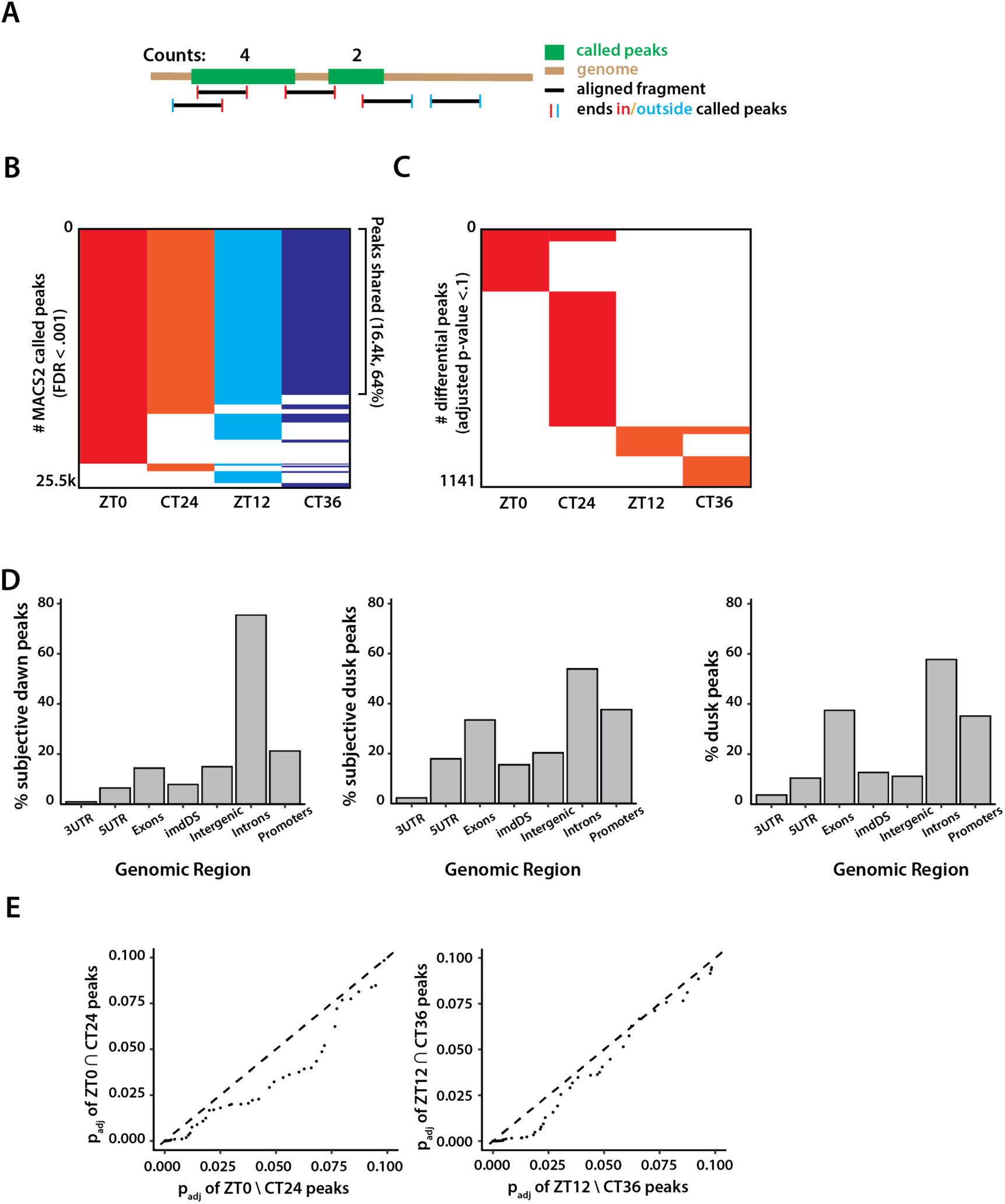
Schema of ATAC signal differential analysis. (**A**) Illustration of how we counted ATAC signal from MACS2 called peaks using the custom edgeCounter package (see Methods). (**B,C**) Binary heatmap comparing MACS2 called peaks under LD and DD conditions (B) and differentially accessible peaks at dawn, dusk, subjective dawn, and subjective dusk (C). (**D**) Distribution of subjective dawn/dusk-accessible peaks with respect to known genomic features reported by ChIPpeakAnno. Distribution of dusk-accessible peaks are also shown (right panel). (**E**) Quantile-quantile plots showing correlation between ZT and CT peaks at dawn (ZT0/CT24) and dusk (ZT12/CT36). Differential peaks that are conserved between ZT and CT conditions (y-axis) shows lower adjusted p-values compared to peaks that are not conserved (i.e., only found in ZT, x-axis).

**Supplementary Figure S4.**
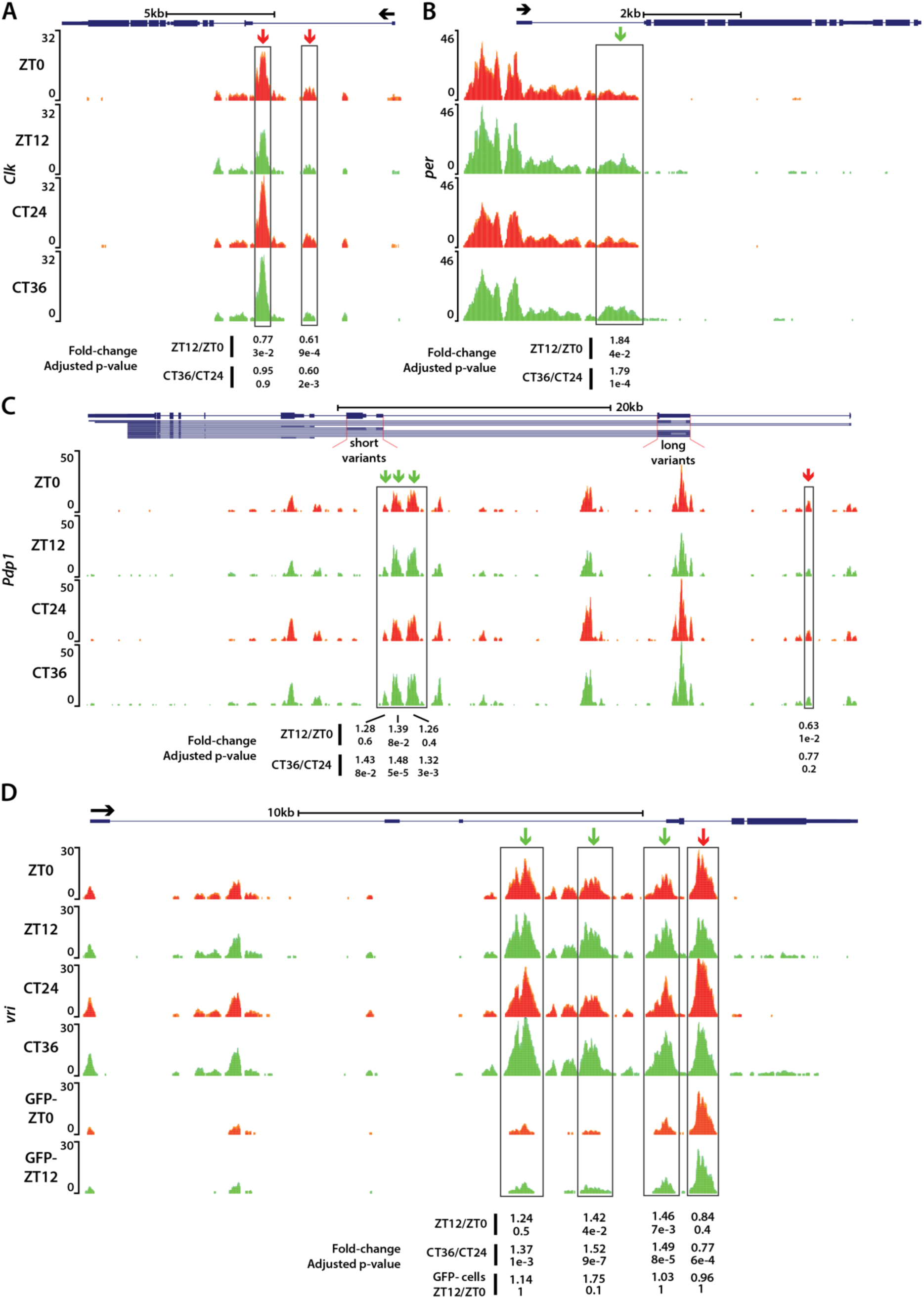
ATAC signal pile-up tracks of core clock genes *Clk, Pdp1 and vri* under LD and DD conditions. (**A,B**) *Clk* locus is more accessible at dawn (ZT0/CT24) compared to dusk while *per* locus shows the opposite pattern. (**C**) Transcript variants of Pdp1 exhibit varying accessibility changes. The “long variants” possess a regulatory element that is more accessible at dusk, whereas the “short variants” display the opposite pattern. (**D**) *vri* locus also shows complex differential peak patterns. It possesses regulatory elements that are more accessible at dawn and dusk in GFP-positive clock neurons. Additionally, *vri* locus also possesses a regulatory element that is more accessible specifically at dusk even in non-clock neurons (GFP-negative). Fold change and adjusted p-value are shown at the bottom of each of the differential peaks identified.

**Supplementary Figure S5.**
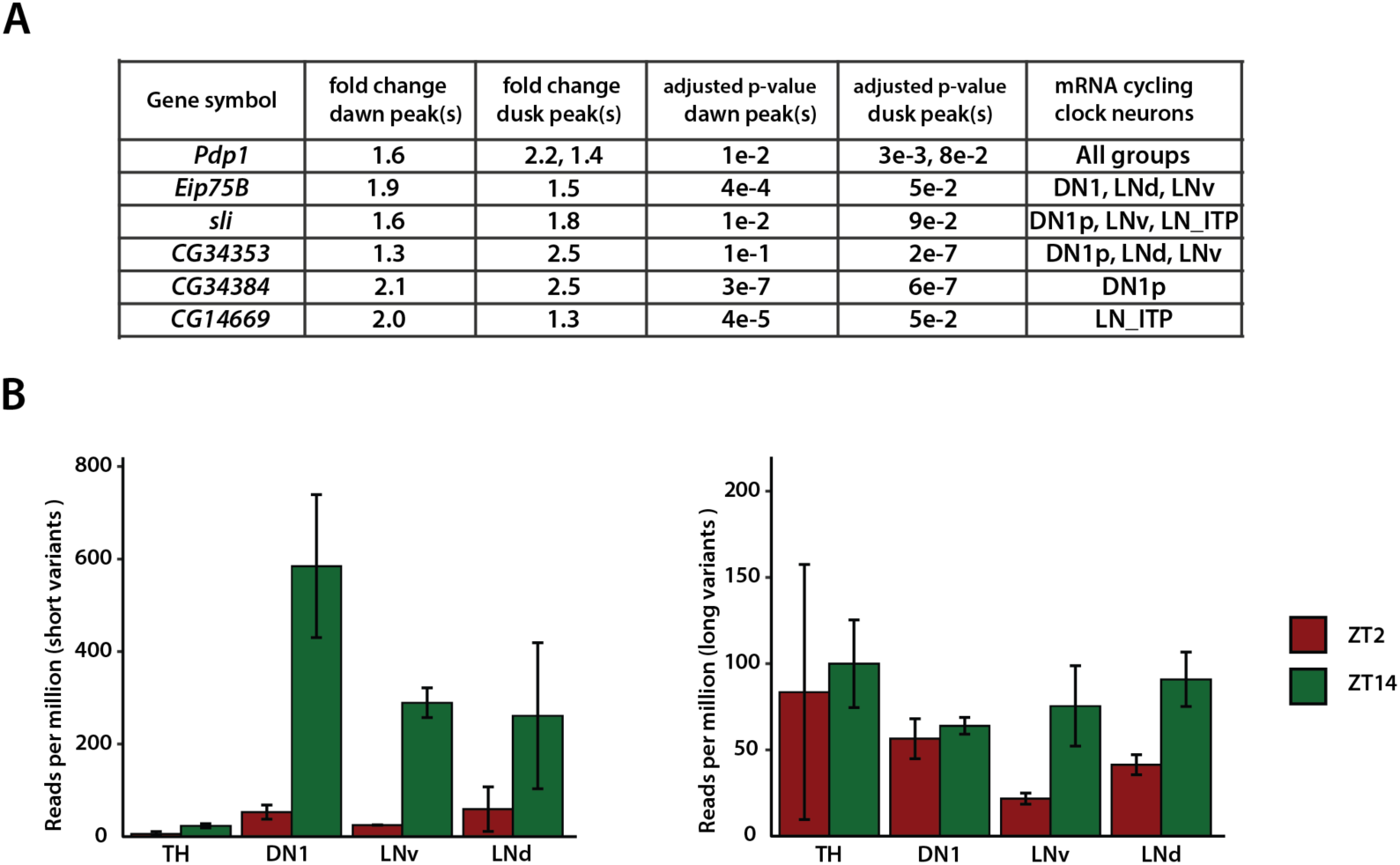
Genes with peaks that are more accessible at both dusk and dawn. (A) List of genes that have differentially accessible peaks at both dusk and dawn. In all, 11 genes meet this criterion, and only those that exhibit cyclical changes in mRNA levels are presented here (6 out of 11). (**B**) Expression level of “short” and “long” transcript variants of Pdp1 in differential clock neuron subgroups (DN1, LNv, LNd) and a non-clock neuron control (dopamine neurons, TH). While short variants show robust cycling profiles in all clock neuron groups, long variants are not cycling in DN1 neurons. Analysis was performed with publicly available data (21).

**Supplementary Figure S6.**
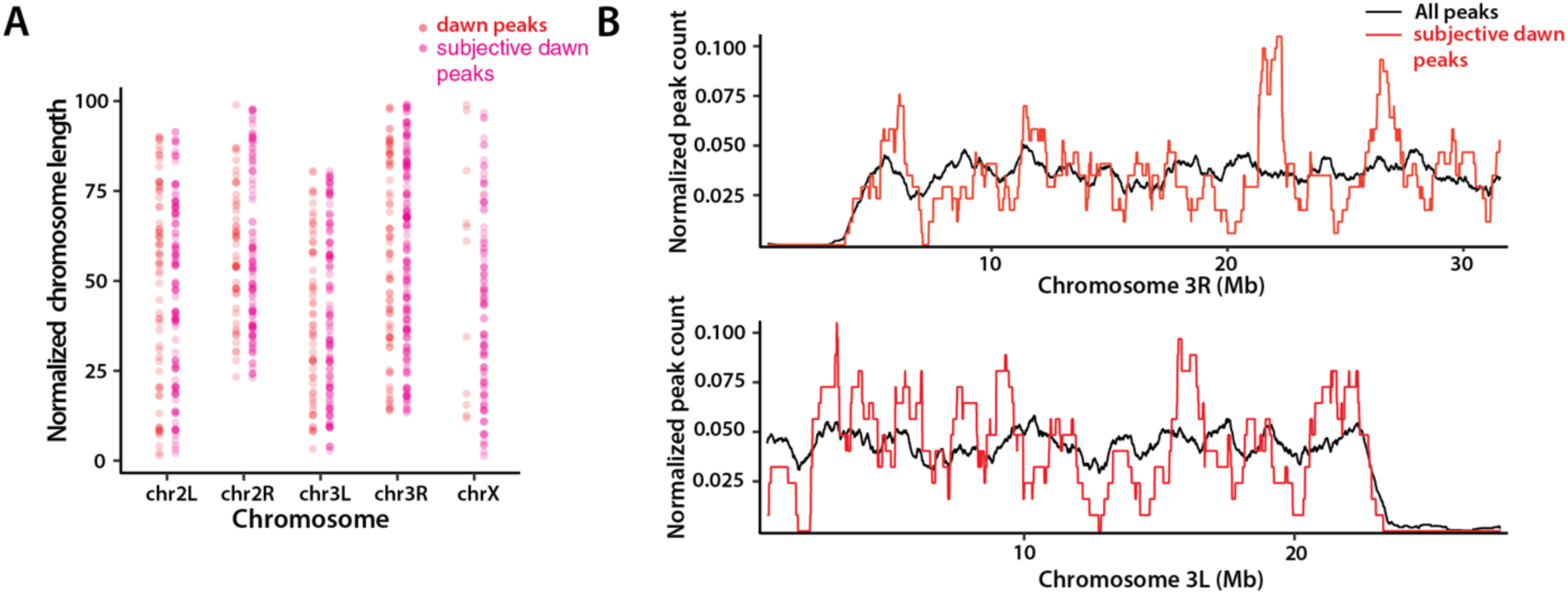
Chromosomal distribution of ATAC peaks that are more accessible at dawn. (A) Distribution of ATAC peaks that are more accessible at dawn (ZT0/CT24) along 1D-chromosome coordinates. Coordinates are normalized by length of the chromosomes (y-axis). (B) Zoomed-in distribution of ATAC peaks that are more accessible at subjective dawn (CT24) along chromosomal arms 3L and 3R.

